# Robust archaeal and bacterial communities inhabit shallow subsurface sediments of the Bonneville Salt Flats

**DOI:** 10.1101/553032

**Authors:** Julia M. McGonigle, Jeremiah A. Bernau, Brenda B. Bowen, William J. Brazelton

## Abstract

We report the first census of natural microbial communities of the Bonneville Salt Flats (BSF), a perennial salt pan at the Utah–Nevada border. Environmental DNA sequencing of archaeal and bacterial 16S rRNA genes was conducted on samples from multiple evaporite sediment layers of the surface salt crust. Our results show that at the time of sampling (September 2016), BSF hosted a robust microbial community dominated by diverse Halobacteriaceae and *Salinibacter* species. Desulfuromonadales from GR-WP33-58 are also abundant in all samples. We identified taxonomic groups enriched in each layer of the salt crust sediment and revealed that the upper gypsum sediment layer found immediately under the uppermost surface halite contains a robust microbial community. We found an increased presence of Thermoplasmatales, Nanohaloarchaeota, Woesearchaeota, Acetothermia, Halanaerobium, Parcubacteria, Planctomycetes, Clostridia, Gemmatimonadetes, Marinilabiaceae and other Bacteroidetes in this upper gypsum layer. This study provides insight into the diversity, spatial heterogeneity, and geologic context of a surprisingly complex microbial ecosystem within this macroscopically-sterile landscape.

**IMPORTANCE:** Over the last ∼13,000 years the Pleistocene Lake Bonneville, which covered a large portion of Utah, drained and desiccated leaving behind the Bonneville Salt Flats (BSF). Today BSF is famous for its use as a speedway, which has hosted many land-speed records and a community that greatly values this salty landscape. Additionally, the salts that saturate BSF basin are extracted and sold as an additive for agricultural fertilizers. The salt crust is a well-known recreational and economic commodity, but the roles of microbes in the formation and maintenance of the salt crust are generally unknown. This study is the first geospatial analysis of microbial diversity at this site using cultivation-independent environmental DNA sequencing methods. Identification of the microbes present within this unique, dynamic, and valued sedimentary evaporite environment is an important step toward understanding the potential consequences of perturbations to the microbial ecology on the surrounding landscape and ecosystem.

## INTRODUCTION

The Great Salt Lake and Bonneville Salt Flats (BSF) are both remnants of Pleistocene Lake Bonneville which drained at the end of the last glacial maximum and desiccated over the last ∼13,000 years (Crittenden, 1963). BSF is famous for its use as a speedway through the last century, which has hosted many land-speed records and a community that greatly values this salty landscape (Noeth, 2002). Over this same time period, the salts that saturate BSF basin have been extracted as an economic commodity, particularly as potash (e.g. KCl) that is sold as an additive for agricultural fertilizers. The hydrology and sedimentology of BSF have been studied periodically throughout the 20^th^ century in relation to potash mining and concerns about the impacts of land use on the environment (Associates, 1979; Brooks, 1991; Eardley, 1962; Kaliser, 1967; L. J. Turk, S.N. Davis, 1973; Lines, 1979; Turk, 1970; White III and Terrazas, n.d.). Human land use has altered many aspects of the hydrology and morphology of the environment that facilitated deposition of the ∼2m thick evaporite salt crust that caps BSF, and the amount and extent of salt present at the site have been observed to change through time (Bowen, B.B., Bernau, J., Kipnis, E.L., Lerback, J., Wetterlin, L. and Kleba, 2018; Bowen et al., 2018).

Examination of modern and ancient salt pan deposits demonstrates that these environments undergo repeated cycles of desiccation (dry saline pan), flooding (brackish lake), and evaporative concentration (drying to dry saline pan) (Bowen and Benison, 2009; Lowenstein, T.K. and Hardie, 1985). While the desiccation stage is most common, it is repeatedly interrupted by flooding events related to individual storms, wet seasons, and spring thaws driving runoff from surrounding mountains. The environmental parameters that influence the hydrology and timing of flooding, evaporation, and desiccation cycles are dynamic (Bowen et al., 2017). For this study, multiple individual strata from several sites spanning the salt crust were sampled at BSF during an extensive sampling campaign to measure the overall stratigraphy and volume of the salt crust (Bowen et al., 2018). The samples were collected in early September 2016 near the end of a three-month long desiccation period.

Microbial communities provide critical ecosystem services by cycling nutrients and providing a biological foundation upon which other organisms can establish themselves. In environments considered “extreme” by human standards such as BSF, microbes are often the only life forms that can survive. The study of microbial life in these ecosystems can elucidate how extremophilic organisms have evolved unique attributes, strategies, and metabolic capabilities that enable them to thrive where most other organisms cannot (Ma et al., 2010; Ventosa et al., 2015). Extreme hypersaline ecosystems select for organisms capable of not only tolerating, but actually requiring high salt for growth (Litchfield and Gillevet, 2002). Microbes in these ecosystems employ either a “salt-in” or “organic-solutes-in” strategy to overcome the problems associated with osmoregulation in a high salt environment (Oren, 1999). In addition, halophiles found in these environments produce enzymes with potential biotechnological, bioremediation, and medical applications (Ventosa and Nieto, 1995).

The role of microbes in BSF salt crust processes are generally unknown. Identification of the microbes present within this unique, dynamic, and valued sedimentary evaporite environment is an important step toward understanding the potential consequences of perturbations to the microbial ecology on the surrounding landscape and ecosystem. This study is the first geospatial analysis of microbial diversity at BSF using cultivation-independent environmental DNA sequencing methods. This study provides insight into the diversity, spatial heterogeneity, and geologic context of a surprisingly complex microbial ecosystem within this macroscopically-sterile landscape.

## METHODS

### Sampling Location and Collection

Samples were collected from eight pits dug with sterile tools in September 2016 **(*Table S1, Fig. S1*)**. Distinct sediment layers visually identified by color and textural changes were sampled at each site. Samples were immediately placed on ice and transported within 12 hours to the University of Utah where they were stored at −80 °C until microbiological analyses were performed in the following 6 months.

### Elemental and Sedimentological Analysis

A portion of each homogenized frozen sample was separated and analyzed for major and trace elements by ActLabs. Major and trace elements were analyzed via Aqua Regia ICP (Inductively Coupled Plasma; method 1E3). Anions were analyzed via Ion Chromatography (method 6B). Using this elemental data along with visual characteristics, each sample was grouped into one of five sediment categories. Elemental concentrations were used to reconstruct relative weight mineralogy based upon stoichiometry of gypsum, halite, and clay. Elemental concentrations beyond detection limits were derived using this method.

In addition to these samples, additional blocks of representational material were collected at the same time of sampling from sites 56, 35, and 12B. These blocks were processed by Wagner Petrographic to create petrographic thin sections for visual microscopy analysis. Blocks were desiccated to remove any pore waters, impregnated with clear or colored epoxy. Several thin sections were made from each block by cutting 50 by 75 mm blocks and mounting them on glass slides. Samples were cut to thicknesses ranging from 0.5 to 2 mm. The samples were then analyzed at the University of Utah using an Olympus BX53M microscope with 1000x magnification and a Zeiss M2 Petroscopic Microscope with an Axiocam Camera and Zen 2 Pro software.

### DNA Extraction and Sequencing

DNA was extracted from salt crust samples using a protocol from Brazelton et al. (2010) modified for extremely high-salt material (Brazelton et al., 2010). The modified protocol is available on our lab’s website (https://baas-becking.biology.utah.edu/data/category/18-protocols) and summarized here. Sediment samples were crushed and homogenized with a sterile mortar and pestle, and 0.25 g subsamples were placed in a DNA extraction buffer containing 0.1 M Tris, 0.1 M EDTA, 0.1 M KH_2_PO_4_, 0.8 M guanidium HCl, and 0.5 % Triton-X 100. For lysis, samples were subjected to one freeze-thaw cycle, incubation at 65 °C for 30 minutes, and to beating with 0.1 mm glass beads in a Mini-Beadbeater-16 (Biospec Products). Purification was performed via extraction with phenol-chloroform-isoamyl alcohol, precipitation in ethanol, washing in Amicon 30K Ultra Centrifugal filters, and final clean-up with 2x SPRI beads (Rohland and Reich, 2012). DNA quantification was performed with a Qubit fluorometer (ThermoFisher).

Archaeal and bacterial 16S rRNA gene amplicon sequencing was conducted by the Michigan State University genomics core facility. The V4 region of the bacterial 16S rRNA gene was amplified with dual-indexed Illumina fusion primers with the 515F/806R primers as described by Kozich *et al.* (Kozich et al., 2013). The V4 region of the archaeal 16S rRNA gene was amplified with the A519F/Arch958R primers (Klindworth et al., 2013). Amplicon concentrations were normalized and pooled using an Invitrogen SequalPrep DNA Normalization Plate. After library QC and quantitation, the pool was loaded on an Illumina MiSeq v2 flow cell and sequenced using a standard 500 cycle reagent kit. Base calling was performed by Illumina Real Time Analysis (RTA) software v1.18.54. Output of RTA was demultiplexed and converted to fastq files using Illumina Bcl2fastq v1.8.4.

### Analysis of 16S rRNA Amplicon Data

Quality screening and processing of all 16S rRNA amplicon sequences was conducted with the DADA2 R package (Callahan et al., 2016). Using this package primer contaminants and chimeras were removed, reads were trimmed and filtered based on quality, and sequence variants likely to be derived by sequencing error were identified. The final amplicon sequence variants (ASVs) are considered to be true variants and are analyzed in the same method as traditional operational taxonomic units (OTUs). Taxonomic classification of all ASVs was performed with the DADA2 package and the SILVA reference database (**NRv123**).

Sequence counts for each ASV were transformed with a biomass correction by multiplying the proportion of each ASV’s contribution to each sample’s total sequence count by the estimated number of total cells in that sample. Total cells were estimated by the total ng of extracted DNA (as measured by the Qubit fluorometer) and an assumption of 2×10^−6^ ng of DNA per cell. All downstream analyses were performed with these biomass-weighted relative abundances. Differential abundance of ASVs in each layer was performed using edgeR (McMurdie and Holmes, 2013a) and phyloseq (McMurdie and Holmes, 2013b). An ASV was considered to be significantly enriched in a layer if its differential abundance passed a false discovery rate threshold of 0.05. Multivariate analysis was done using phyloseq (McMurdie and Holmes, 2013b) and Bray-Curtis and Simpson indices were calculated using vegan (Jari Oksanen, F. Guillaume Blanchet, Michael Friendly, Roeland Kindt, Pierre Legendre, Dan McGlinn, Peter R. Minchin, R. B. O’Hara, Gavin L. Simpson, Peter Solymos, M. Henry H. Stevens, 2018).

### Accession numbers

All sequence data are publicly available at the NCBI Sequence Read Archive under BioProject PRJNA522308. All SRA metadata, supplementary material, and protocols are archived with the following DOI: http://doi.org/10.5281/zenodo.2564239. All custom software and scripts are available at https://github.com/Brazelton-Lab.

## RESULTS

### Elemental and Sedimentological Analysis

Thin sections and reconstructed mineralogy from elemental data were used to group each sample into one of five classifications: Group 1 (surface halite), Group 2 (upper gypsum), Group 3 (lower halite), Group 4 (lower gypsum), and Group 5 (halite mixed with gypsum).

Thin section and elemental analysis and microscopy reveal different layers of distinct sedimentological morphology (***Fig. 1*, *Fig. S2***). Primary minerals were identified as halite (NaCl), gypsum (CaSO_4_*2H_2_O), and clay minerals (Fe, Mg, and Al). The majority of samples were above reporting limits for sodium. All minerals are reported as weight % of composition (***Table S1, Figs. S3-S5***). The relative enrichment of select major and trace elements was determined for each sediment classification (***Table 1***). Groups with mean values above the cutoff limit were reported as enriched with an element.

**Figure 1:**
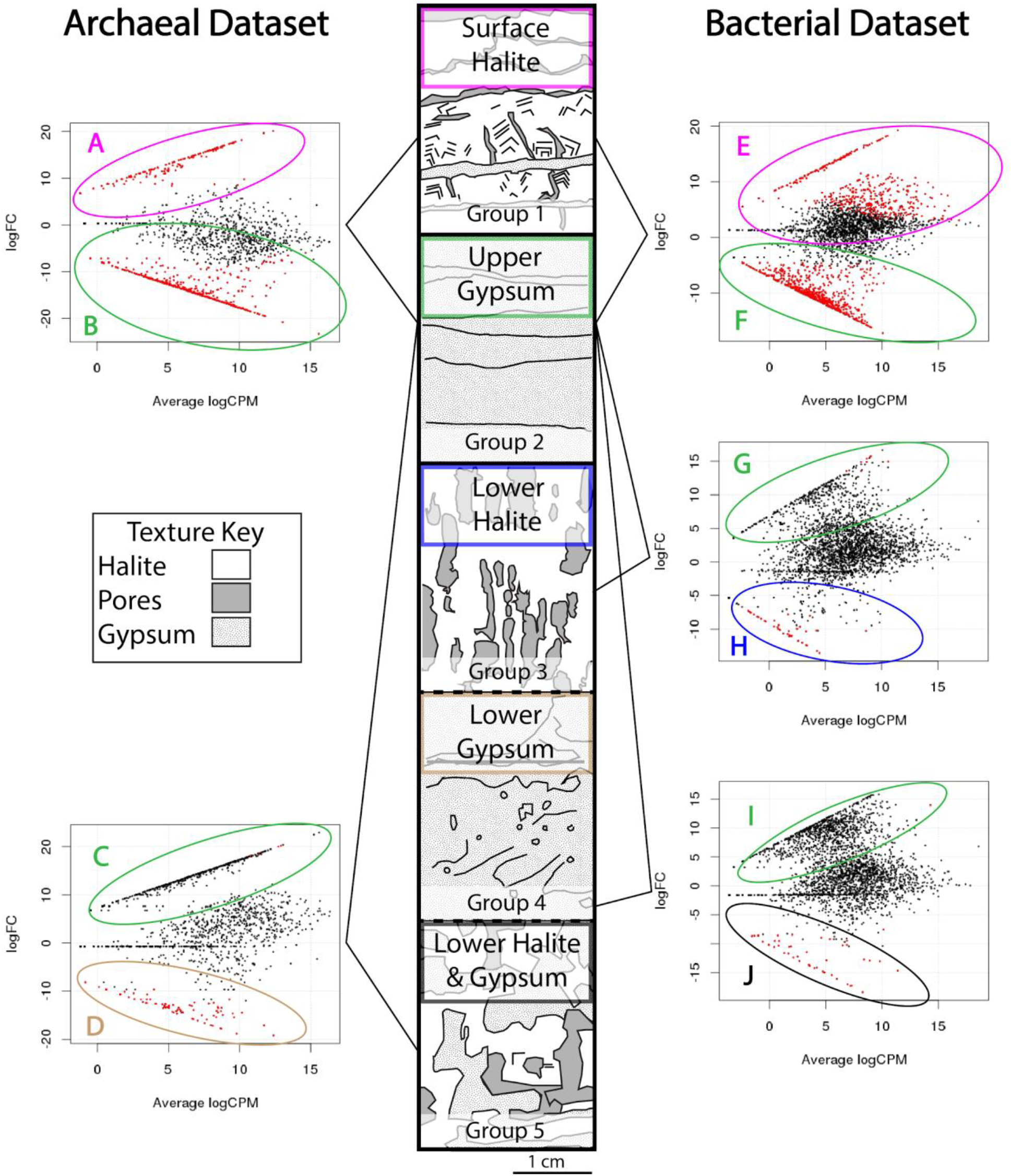
Sedimentology and differential abundance results (EdgeR) for archaeal and bacterial datasets. Red dots indicate an ASV with a significantly higher abundance (false discovery rate <0.05) in the indicated category. Representative stratigraphic column presented here. Dashed lines represent stratigraphic shifts that only occurred at some sites. General Stratigraphy consists of groups 1-5. Group 1, surface halite, contains efflorescent halite and halite crystals with abundant fluid inclusions. Group 2, upper gypsum, consists of gypsum grains with varying proportions of halite, this group has the highest proportion of clays. Group 3, lower halite, has the highest proportion of pore space and is mineralogically similar to group 1. Group 4, lower gypsum, contains coarser gypsum grains than group 2, this group also smelled of sulfur. Group 5, lower halite and gypsum, forms when pores in lower halite fill with gypsum.

**Table 1:** Sample means for sediment categories. Bold values indicate element is enriched in samples.

|  | Cutoff Value | Surface Halite (G1) | Upper Gypsum (G2) | Lower Halite (G3) | Lower Gypsum (G4) | Halite/Gypsum (G5) |
| --- | --- | --- | --- | --- | --- | --- |
| **Mn** | 15 mg/L | 10.25 | **38** | 5.2 | **19.5** | 10.5 |
| **Zn** | 10 mg/L | **47.5** | **80.29** | **41.2** | 5 | **19** |
| **Al** | 0.10% | 0.01 | **0.14** | 0.02 | **0.1** | 0.05 |
| **Ba** | 20 mg/L | 1.75 | **29.29** | 4.6 | **22.5** | **22** |
| **Ca** | 5% | 1.99 | **7.97** | 3.19 | **7.92** | **6.93** |
| **Fe** | 0.05% | 0.02 | **0.13** | 0.02 | **0.09** | 0.04 |
| **K** | 0.15% | **0.2** | **0.21** | 0.09 | **0.17** | **0.15** |
| **Mg** | 0.20% | 0.13 | **0.4** | 0.07 | **0.25** | 0.16 |
| **S** | 2% | 1.54 | **5.67** | 2.6 | **6.75** | **5.86** |
| **Sr** | 200 mg/L | 106.38 | **505.29** | 146.2 | **465** | **472** |
| **Cl** | 450,000 mg/L | **541125** | 139700 | **520800** | 201750 | 384500 |

Thin sections revealed Group 1 (surface halite) consists of puffy fine efflorescent halite crystals followed by larger halite crystals (up to 0.5 cm wide) with fluid inclusions. Group 1 samples are composed of predominantly halite with 2.5-27% gypsum and trace amounts (<0.5%) of clay. Group 1 is relatively enriched in chloride, potassium, zinc, and lead (from most to least abundant).

Group 2 (upper gypsum) consists of layered fine to medium sized gypsum grains. Group 2 samples contain 29-39% gypsum and 6-22% halite. Samples 56-2 and 35-2 are mineralogical outliers in this group as they have higher amounts of halite. Thin sections from sites 35 and 56 show relatively thin (∼5-20 mm thick) layers of fine-grained gypsum adjoined by halite (***Fig. S2***). Group 2 samples contain the highest amount of clay minerals of any strata at 0.3-1.7%. Reconstructed mineralogy only accounted for 41-61% of Group 2’s mass. Group 2 is enriched in sulfur, calcium, magnesium, aluminum, potassium, iron, strontium, zinc, lead, and manganese.

Group 3 (lower halite) samples are cemented halite with porous vertical dissolution pipes running through it. These pipes are typically 10mm – 30mm wide, but some are > 1 cm. Elemental data from group 3 is similar to Group 1 (surface halite). Group 3 is enriched in chloride, and zinc.

Group 4 (lower gypsum) samples contain coarse black organic or clay coated gypsum grains (medium to coarse, some very coarse) that smell of sulfur. Samples in this category have similar amounts of gypsum (32.5-38%) but differing amounts of halite (6.5-54%). Group 4 samples are enriched in sulfur, calcium, magnesium, aluminum, potassium, iron, strontium, and barium.

Group 5 (halite mixed with gypsum) consists of a more chaotic halite (53-61%) supported framework made from vestigial dissolution pipes filled by pore space and 28-33% gypsum. Group 5 is enriched in chloride, calcium, sulfur, strontium, and barium.

### Bacterial and Archaeal Community Composition

All sediment samples produced 1,845,510 merged paired reads with the bacterial primer set and 552,906 merged paired reads with the archaeal primer set. From these, DADA2 identified 2,740 amplicon sequence variants (ASVs) in the bacterial dataset and 1,646 ASVs in the archaeal dataset. DNA extraction yields and ASV counts per sample are shown in ***Table S2***. These ASV counts were transformed with the DNA extraction yields with the procedure described in the methods to obtain biomass-weighted environmental abundances for downstream analyses. At most sites, samples in Group 1 and Group 2 have higher DNA yields and ASV counts than samples from Group 3, 4, and 5. Samples from Group 1 and 2 sequenced more successfully than those in other groups, particularly with archaeal primers.

There were 1,552 ASVs (56% of the total ASVs) in the bacterial dataset that were assigned archaeal taxonomy. The 515F/806R bacterial primers are known to amplify both bacteria and archaea, so we chose to include the archaeal ASVs in the bacterial dataset rather than remove such a large proportion of the total sequence diversity (Apprill et al., 2015; Parada et al., 2016; Walters et al., 2015). Most archaeal ASVs in the bacterial dataset overlap classifications found in the archaeal dataset, but the archaeal dataset generally provided more specific classifications, particularly for the class Thermoplasmata. The exceptions are Woesearcheota and Nanohaloarchaeota, which are only found in the bacterial dataset.

All samples in the bacterial dataset are dominated by diverse Halobacteriaceae and *Salinibacter* species (***Fig. S6***). All samples in the archaeal dataset are dominated by Halobacteriaceae from 27 different classified genera (***Fig. S7***). Desulfuromonadales from GR-WP33-58 are also abundant in all samples. Less dominant taxa generally increase in abundance with depth at most sites. The most dominant cyanobacteria are classified as *Lyngbya* and are more abundant in upper surface halites. Nanohaloarchaeota and ASVs classified to the Soil Crenarchaeotic Group also appear more abundant in these upper sediments. Woesearchaeota, Archaeoglobaceae, Aenigmarchaeota (DSEG), and Methanomicrobia (ST-12K10A) appear more abundant in lower layers. In the bacterial dataset, the relative abundances of Acetothermia, Rhodovibrio, Marinilabiaceae, Desulfovermiculus, Gemmatimonadetes, Phycisphaeraceae, Thiohalorhabdus, and Ectothiorhodospiraceae are greater in lower layers. In both datasets, the relative abundances of ASVs classified to Thermoplasmata are greatest in the middle layer.

Multivariate analysis of beta diversity for the bacterial and archaeal datasets did not reveal any significant correlations between community compositions and location or sediment depth. (***Fig. S8, Fig. S9***). Similarly, alpha diversity, as measured by the Simpson index, did not exhibit any consistent patterns with location or sediment depth (***Table S3***). These results highlight the general lack of significant variation among the whole-community compositions prior to differential abundance analyses.

### Differential Abundance

Despite the lack of variation at a whole community level, we investigated whether specific ASVs are significantly enriched in one or more sediment layers. Differential abundances of ASVs were calculated edgeR (McMurdie and Holmes, 2013a) by comparing each of the sediment groups to each other (***Fig. 1***). For the bacterial dataset, comparisons were done between: Group 1-2, Group 2-3, and Group 2-4. For the archaeal dataset, comparisons were done between: Group 1-2 and Group 2-5. All other comparisons were not possible due to lack of data from particular samples.

In the bacterial dataset, comparisons between Group 1 and 2 identified 429 ASVs with significantly greater abundance (determined by a false discovery rate of <0.05) in Group 1 (surface halite) and 660 ASVs with significantly greater abundance in Group 2 (upper gypsum) (***Fig. 2*, *Fig. S10***). Diverse Halobacteriaceae make up 37% of the surface halite-enriched ASVs. *Salinibacter* species comprise another 43% of these ASVs. The remaining ASVs enriched in Group 1 are related to the cyanobacteria *Euhalothece*, Desulfuromonadales, Nanohaloarchaeota, and unclassified members of Chitinophagaceae. The enriched taxa become more diverse as the sediment transitions from surface halite to the upper gypsum layer. Taxa with a greater abundance in these Group 2 sediments include Thermoplasmatales, Nanohaloarchaeota, Woesearchaeota, Acetothermia, Halanaerobium, Parcubacteria, Planctomycetes, Clostridia, Gemmatimonadetes, Marinilabiaceae and other Bacteroidetes. Unclassified bacterial and archaeal ASVs are also more abundant in Group 2.

**Figure 2:**
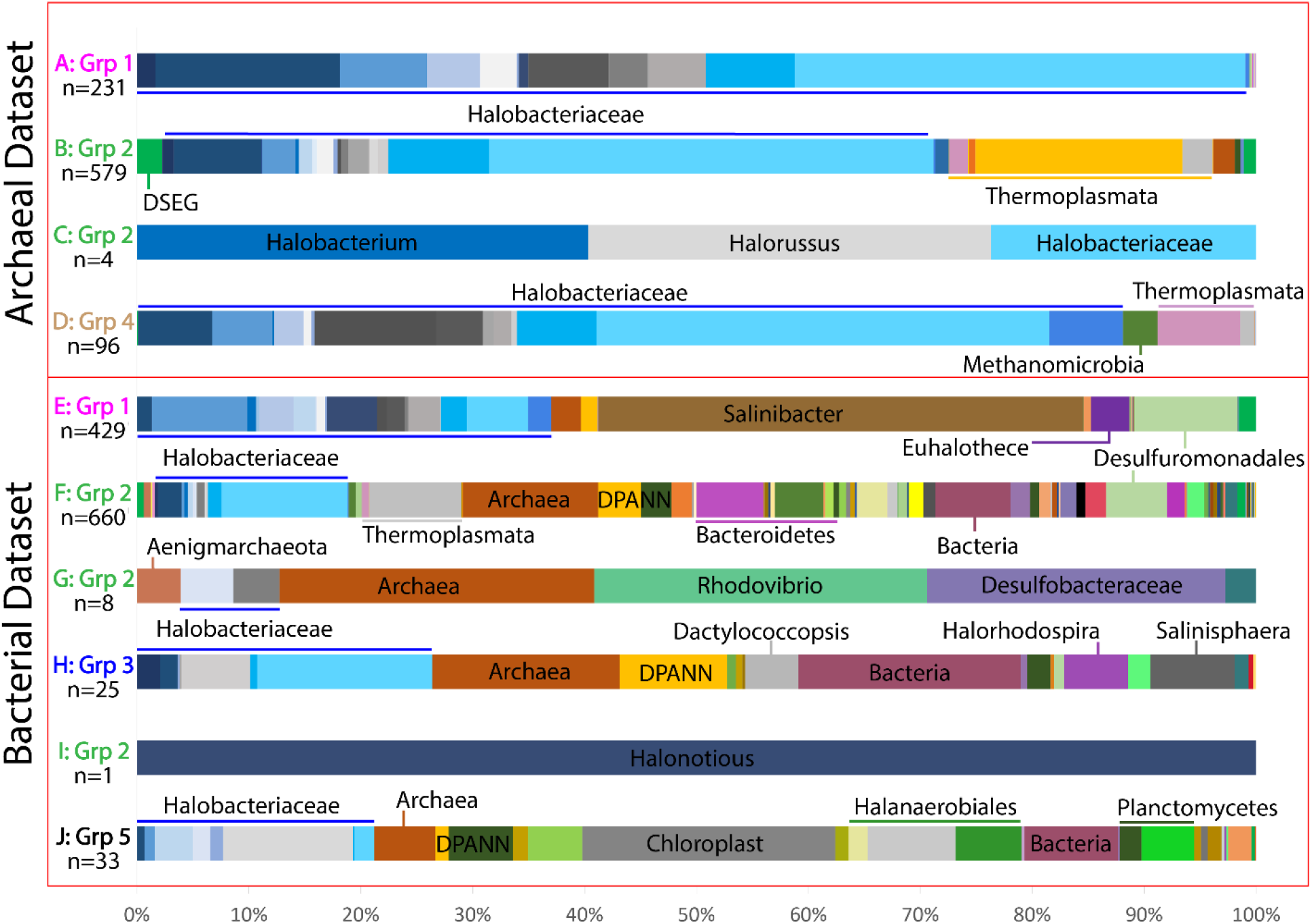
Abundance of ASVs enriched in each group as determined by differential abundance (EdgeR) comparisons shown in Figure 1. Number of ASVs making up the total plot are indicated below the comparison ID. A full legend can be found in Fig. S11.

In the archaeal dataset, comparisons between Group 1 and 2 identified 231 ASVs with significantly higher abundance in Group 1 (surface halite) and 579 ASVs with significantly higher abundance in Group 2 (upper gypsum). Diverse Halobacteriaceae make up 99% of the surface halite-enriched ASVs, with the majority of these belonging to unclassified species. The remaining 1% belong to members of DSEG and Thaumarchaeota. The ASVs enriched in Group 2 are also dominated by diverse Halobacteriaceae as well as Thermoplasmatales. A few ASVs enriched in Group 2 belong to DSEG and unclassified Archaea.

The number of enriched ASVs identified by differential abundance comparisons between upper gypsum and deeper sediment layers are much lower. In the bacterial dataset, comparing Group 2 (upper gypsum) and 3 (lower halite) revealed 8 enriched ASVs in Group 2, mostly from Aenigmarchaeota, Rhodovibrio, and Desulfobacteraceae, and 25 Group 3-enriched ASVs heavily represented by Halobacteriaceae.

In the bacterial dataset, comparison of Group 2 (upper gypsum) and 4 (lower gypsum) identified only one differentially abundant ASV in Group 2 samples, which was classified as *Halonotius*, a species of Halobacteriaceae. 33 ASVs were identified as enriched in Group 4, including diverse Halobacteriaceae, DPANN (Nanohaloarchaeota and Woesearchaeota), candidate division SR1, chloroplasts, Halobacteroidaceae, and Clostridia such as Halanaerobium.

In the archaeal dataset, comparisons between Group 2 (upper gypsum) and 5 (halite mixed with gypsum) found only 4 ASVs that were more abundant in Group 2 samples, all belonging to Halobacteriaceae. There were 96 ASVs enriched in Group 5 samples. These ASVs were classified as Aenigmarchaeota, diverse Halobacteriaceae, Methanomicrobia, Thermoplasmata, and Thaumarchaeota.

## DISSCUSION

### The Bonneville Salt Flats hosts a robust microbial ecosystem

Despite its barren appearance on the surface, the Bonneville Salt Flats (BSF) appears to harbor a robust microbial ecosystem featuring a range of metabolic capabilities potentially including phototrophy, sulfur and nitrogen cycling metabolisms, and heterotrophy. Although hypersaline systems can be harsh environments for life, they have been previously reported to contain high biomass (Çınar and Mutlu, 2016; Litchfield and Gillevet, 2002; Maturrano et al., 2006). Our results indicate that BSF is no exception. If we assume 2×10^−6^ ng of DNA per cell, we estimate 10^7^ – 10^9^ cells per gram of sample from our quantifications of environmental DNA. This is likely to be an over-estimate, since it is known that salt can preserve DNA, and our environmental DNA could include a significant extracellular fraction (Borin et al., 2008; Rhodes et al., 2011). Nevertheless, the low range of our biomass estimate is consistent with other reported values around 10^6^- 10^7^ cells mL^−1^ (Çınar and Mutlu, 2016; Litchfield and Gillevet, 2002; Maturrano et al., 2006).

The surface halite and upper gypsum sediment layers (Groups 1 and 2) have higher DNA yields than deeper samples (***Table S2***). These upper sediments have availability to sunlight and oxygen while still retaining protection from UV radiation within the evaporite crusts. Additionally, sulfur minerals in the upper gypsum layer likely provide an energy source for chemotrophic microbes living in these upper layers. Therefore, the upper gypsum layer is likely to have more abundant and more active phototrophic and chemotrophic microbial communities compared to deeper layers.

### A microbial community shift occurs along a mineralogical shift in the sediment

Although the depth of each sediment layer varied per site, microbial communities seemed homogenous among samples within each sediment category. For example, at site 12B, the surface halite was only 3 mm, while at site 56 the surface halite was 25mm (***Fig. S11, Table S1***). In spite of these differences in depth, these surface halite samples hosted remarkably similar archaeal and bacterial communities (***Fig. S6, Fig. S7***). This trend is still evident even between the two samples with the greatest spread between sampling depths, site 12B where the upper gypsum layer was sampled at a depth of 3mm-7mm and site 41 where the gypsum layer was sampled at a depth of 25mm – 90mm. Both of these samples appear similar in mineral classification and relative abundance of microbial taxa. These results suggest microbial communities shift along mineralogical rather than depth gradients at BSF.

Upper halite crusts (Group 1) are dominated by heterotrophs from *Salinibacter* and Halobacteriaceae. Halobacteriaceae gain most of their energy through heterotrophy but are also capable of harnessing sunlight for supplemental ATP production with the specialized pigment bacteriorhodopsin (Hartmann et al., 1980; Sharma et al., 2007; Spudich, 1998). Key primary producers in the top halite layer are likely cyanobacteria, particularly *Euhalothece*, and the salt tolerant algae *Dunaliella salina*. Although we did not perform 18S rRNA sequencing to characterize the eukaryotic community, one ASV has 100% identity to the 16S rRNA gene of the *Dunaliella* chloroplast. This ASV was only absent at 3 sites: 12B, 67B, and 41. Sites 12B and 67B have the highest proportion of gypsum for samples in Group 1. These sites are also located in a region with the thinnest surface halite (***Fig. S11***).

Thin sections show surface halites contain large pore spaces and fluid inclusions within the halite crystals (***Fig. 1***). This upper phototrophic community at BSF likely resides in tandem with heterotrophs in these pore spaces of the halite structure or as a filamentous biofilms attached to the surface of halite crystals (Spear et al., 2003). Microbes in this surface layer may also be encapsulated within fluid inclusions. Members of this community are likely active during the flooding stage of BSF, when upper halite dissolves and brine exists on the surface. They may become encapsulated once again as the brine precipitates into halite crystals during the next desiccation cycle.

Our results indicate that more genera are enriched in the upper gypsum sediments (Group 2) found beneath the surface halite layer than in other types of sediment (***Fig. 2*, *Table 2***). These genera may be concentrated in this layer because gypsum, halite, and other evaporite minerals provide some protection from harsh environmental conditions (such as UV radiation and extreme temperature fluctuations) while still transmitting enough sunlight to support photosynthesis (Friedmann, 1982). The higher surface area of clay minerals present in these samples likely provides a more suitable substrate for growth than the surface halite.

**Table 2:** Number of different Orders (or lowest assigned taxonomic group if not classified to Order) represented in the enriched ASVs from differential abundance comparisons.

|  | Grp 1 | Grp 2 | Grp 3 | Grp 4 | Grp 5 |
| --- | --- | --- | --- | --- | --- |
| # Taxonomic Groups in Bacterial Dataset | 34 | 66 | 19 | 24 | - |
| # Taxonomic Groups in Archaeal Dataset | 10 | 16 | - | - | 6 |

In general, bacterial taxa associated with sulfur metabolism (e.g. *Thiohalorhabdus, Desulfovermiculus*, Desulfobacteraceae, and Desulfobulbaceae) are enriched in the upper gypsum (Group 2) layer. Phototrophs enriched in upper gypsum sediments include members of Ectothiorhodospiraceae, the purple sulfur bacteria. These anaerobic organisms produce sulfur globules outside of their cells. Thermophilic organisms (Thermoplasmatales, Archaeoglobaceae, and Thermosulfurimonas) commonly found in environments containing sulfur compounds are also enriched in this upper gypsum layer. These results are strongly suggestive that the abundant sulfur minerals found in this layer (***Table S1***) are metabolized by the resident microbes. Interestingly, methanogens and acetogens who often compete with sulfate-reducing bacteria, are also enriched in the upper gypsum layer (Kato et al., 2014). Methanogens in salt flats often utilize methylated compounds, rather than carbon dioxide, allowing them to coexist with acetogens and sulfate-reducing bacteria (García-Maldonado et al., 2015).

The lowest layers sampled (Groups 4 and 5) contain chemoheterotrophic (Thermoplasmata and Planctomycetes) and fermentative taxa perhaps dependent on primary production of the phototrophic community in upper layers. Strictly anaerobic fermenters from the class Clostridia (particularly the genus *Orenia* found within the order Halanaerobiales) were enriched in deeper samples where oxygen is likely more depleted. Aerobic heterotrophs from the Proteobacteria, such as *Acinetobacter* and *Sandaracinus*, are also enriched here when compared to higher sediments suggesting oxygen may not be completely depleted at this depth. The dissolution pipes and larger grain sizes found in the lower gypsum, lower halite, and lower halite mixed with gypsum layers may enable more air circulation at these depths (***Fig. S2***).

### The microbial community of BSF is similar to previously described hypersaline ecosystems

Only one culture-independent study of microorganisms at the Bonneville Salt Flats (BSF) has been conducted previously (Lynch, 2015). This study focused on the adjacent basin Pilot Valley but also included one sampling location at the BSF. The genus *Halomina* was the most dominant member of the Halobacteriaceae in this BSF sample. *Halomina* sequences were also identified in all of our samples, but the most abundant Halobacteriaceae genera in our dataset were *Haloarcula* and *Halapricum* (***Fig. S6, Fig. S7***). Culture-dependent studies at BSF have only been successful at isolating *Haloarcula, Halorubrum, Halobacterium*, and *Salicola* species (Boogaerts, 2015; Gary M. King, 2015).

The microbial diversity at BSF is comparable to that of other dry saline environments. Archaeal communities are often dominated by members of Halobacteriaceae (Di Meglio et al., 2016; Kambourova et al., 2017; Stivaletta et al., 2009). Bacterial communities are often dominated by Bacteroidetes (*Salinibacter*) and Proteobacteria (Rasuk et al., 2014; Stivaletta et al., 2011; Vogt et al., 2017). Members from Crenarchaeota, Gemmatimonadetes, Verrucomicrobia, SRB from Deltaproteobacteria, and Clostridia are also commonly reported (Caton and Schneegurt, 2012; Kim et al., 2012; McKay et al., 2016). We found ASVs at BSF that are closely related to all of these commonly reported species (***Fig. S6, Fig. S7***). *Euhalothece* is the most commonly reported dominant Cyanobacteria in hypersaline environments(Caton and Schneegurt, 2012; Lindsay et al., 2017; McKay et al., 2016; Schneider et al., 2013; Spear et al., 2003; Stivaletta et al., 2011; Vogt et al., 2017). Some of our ASVs classified to *Euhalothece*, but we found *Lyngbya* to be the most dominant Cyanobacteria at BSF.

Interestingly, microbial species commonly reported in hypersaline lake environments, such the Great Salt Lake, Lake Chaka, and the Salton Sea are also found in our BSF samples (Almeida-Dalmet et al., 2015; Bowman et al., 2000; Dillon et al., 2013; Jiang et al., 2007; Lindsay et al., 2017; Schneider et al., 2013; Swan et al., 2010), suggesting physiological and/or ecological connections between hypersaline aquatic and sediment systems.

## CONCLUSION

The Bonneville Salt Flats hosts thriving microbial communities dominated by diverse Halobacteriaceae and *Salinibacter* species that are apparently adapted to the challenges of this extreme ecosystem. Many of the archaeal and bacterial taxa are most abundant beneath the halite crust within the upper gypsum sediments, where microbes are likely metabolizing sulfur-containing minerals and can still rely on light penetration for phototrophy.

This study represents the first comprehensive culture-independent characterization of the microbial communities inhabiting the Bonneville Salt Flats, and future work in this understudied ecosystem could address the following questions. Which microbes in this ecosystem are crucial for the cycling of nutrients, such as sulfur and nitrogen? Are the haloarchaea dependent on light filtered through the surface halite crust, and are they involved in surface halite formation? Is there evidence for dispersal and exchange with nearby hypersaline environments such as Pilot Valley and the Great Salt Lake? How does the microbial community shift with seasonal desiccation and flooding stages? What sediment interfaces are microbes inhabiting? Do the compositions of endolithic microbes and microbes preserved in fluid inclusions differ from microbes within sediment interfaces? Has anthropogenic use of this environment for racing and brine mining changed the microbial composition of this area?

Answering these questions would extend our understanding of the presence and preservation of life not only in extreme environments, but in any environment where salt deposits are formed. Halite crystal growth bands can preserve microbial life present in the original air-water and water-sediment interface where the salt formed thousands to hundreds of millions of years ago (Lowenstein, T.K. and Hardie, 1985; Sankaranarayanan et al., 2014). Evaporite ecosystems may even reflect life on Earth prior to the Cambrian explosion (Ventosa et al., 2015). Furthermore, because evaporite deposits on other planetary bodies, like Mars, are evidence of past or present water (Barbieri, R. and Stivaletta, 2012; Barbieri and Stivaletta, 2011; Douglas, 2005; Rothschild, 1990), further advances in the characterization of live and preserved microbes in salt deposits will have implications for the development of extraterrestrial life detection strategies.

## ACKNOWLEDGEMENTS

This research was supported by the National Science Foundation Award 1617473, CNH-L: Adaptation, Mitigation, and Biophysical Feedbacks in the Changing Bonneville Salt Flats. We would like to thank Emily Dart and Betsy Kleba for assistance with sample collection, DNA extraction, and general insight on the BSF system. We would also like to thank Alex Hyer and Christopher Thornton for computational assistance.

